# Characteristics of Interactions at Protein Segments External to Globular Domains in the Protein Data Bank

**DOI:** 10.1101/423087

**Authors:** Kota Kasahara, Shintaro Minami, Yasunori Aizawa

**Affiliations:** College of Life Sciences, Ritsumeikan University, 1-1-1, Noji-higashi, Kusatsu, Shiga 525-8577, Japan; Exploratory Research Center on Life and Living Systems, National Institutes for Natural Sciences, 38 Nishigo-Naka, Myodaiji, Okazaki, Aichi 444-8585, Japan; School of Life Science and Technology, Tokyo Institute of Technology, 4259, Nagatsuda-cho, Midori-ku, Yokohama, Kanagawa 226-8501, Japan

**Keywords:** amino acid propensity, loop region, protein data bank, secondary structure, structural bioinformatics

## Abstract

The principle of three-dimensional protein structure formation is a long-standing conundrum in structural biology. A globular domain of a soluble protein is formed by a network of atomic contacts among amino acid residues, but regions external to globular domains, like loop and linker, often do not have intramolecular contacts with globular domains. Although these regions can play key roles for protein function as interfaces for intermolecular interactions, their nature remains unclear. Here, we termed protein segments external to globular domains as *floating* segments and sought for them in tens of thousands of entries in the Protein Data Bank. As a result, we found that 0.72 % of residues are in floating segments. Regarding secondary structural elements, coil structures are enriched in floating segments, especially for long segments. Interactions with polypeptides and polynucleotides, but not small compounds, are enriched in floating segments. The amino acid preferences of floating segments are similar to those of surface residues, with exceptions; the small side chain amino acids, Gly and Ala, are preferred, and some charged side chains, Arg and His, are disfavored for floating segments compared to surface residues. Our comprehensive characterization of floating segments may provide insights into understanding protein sequence-structure-function relationships.

## INTRODUCTION

Elucidating the principles of the three-dimensional (3D) structure formation of proteins is a long-standing conundrum in the field of structural biology [1]. How a sequence of 20 types of amino acid residues in a polypeptide determines its 3D structure remains largely unclear. Toward illumination of this issue, extensive efforts in structural biology have accumulated a massive amount of structural data for proteins in the Protein Data Bank (PDB) [2]. By taking advantage of the wealth of structural data, the field of so-called “structural bioinformatics”, which tackles extracting biological knowledge from structural databases by using techniques of information science, has arisen [3-6]. Statistical analyses of structural elements in the PDB have provided a bird’s eye view on the characterization of the 3D structures of proteins. For example, statistical analyses revealed amino acid propensities associated with several features, e.g., formation of secondary structural elements [7-10] and loop regions [11]. In addition to secondary structure formation, another key feature to establish protein folding is intramolecular contact between amino acid residues distant in the primary structure; in this paper, we refer to this type of intramolecular contact as *non-local*: contacts between neighboring residues are termed *local contacts*. The statistical analyses of the vicinity of amino acid residues in 3D space have been extensively performed to predict and recognize protein folds [12,13]. For example, the residue-wise contact order, defined as the summation over the distance along the sequence between contacting residues, contains significant information regarding 3D structures (Kinjo and Nishikawa, 2005; Kurt et al., 2008). Propensities of non-local contacts play pivotal roles in establishing folds of globular domains.

On the other hand, proteins also have regions without non-local contacts. A typical example is the linker region, which is a flexible segment linking two globular domains. The structural element termed the loop region also tends to have no or only a few non-local contacts. They are also key elements in the 3D structures of proteins. However, in spite of their importance, the nature of the regions without non-local contacts is not well understood.

Here, we performed statistical analyses on PDB entries to investigate the nature of regions without non-local contacts. Tens of thousands of PDB entries were processed, and two types of regions consisting of consecutive amino acid residues were defined: (i) regions with non-local contacts, and (ii) those without non-local contacts. We refer to these regions as (i) *supported* segments and (ii) *floating* segments, respectively (Fig 1). We aim to characterize the floating segments in proteins. On the basis of non-redundant PDB entries, the frequency of these floating segments was analyzed with many features of the segments, e.g., length, secondary structures, accessible surface area (ASA), and intermolecular interactions. We present that an considerable number of residues are floating in protein structures deposited in the PDB. While amino acid preferences of floating segments are similar to those of exposed residues, some amino acids exhibited a unique propensity for floating segments.

**Figure 1.**
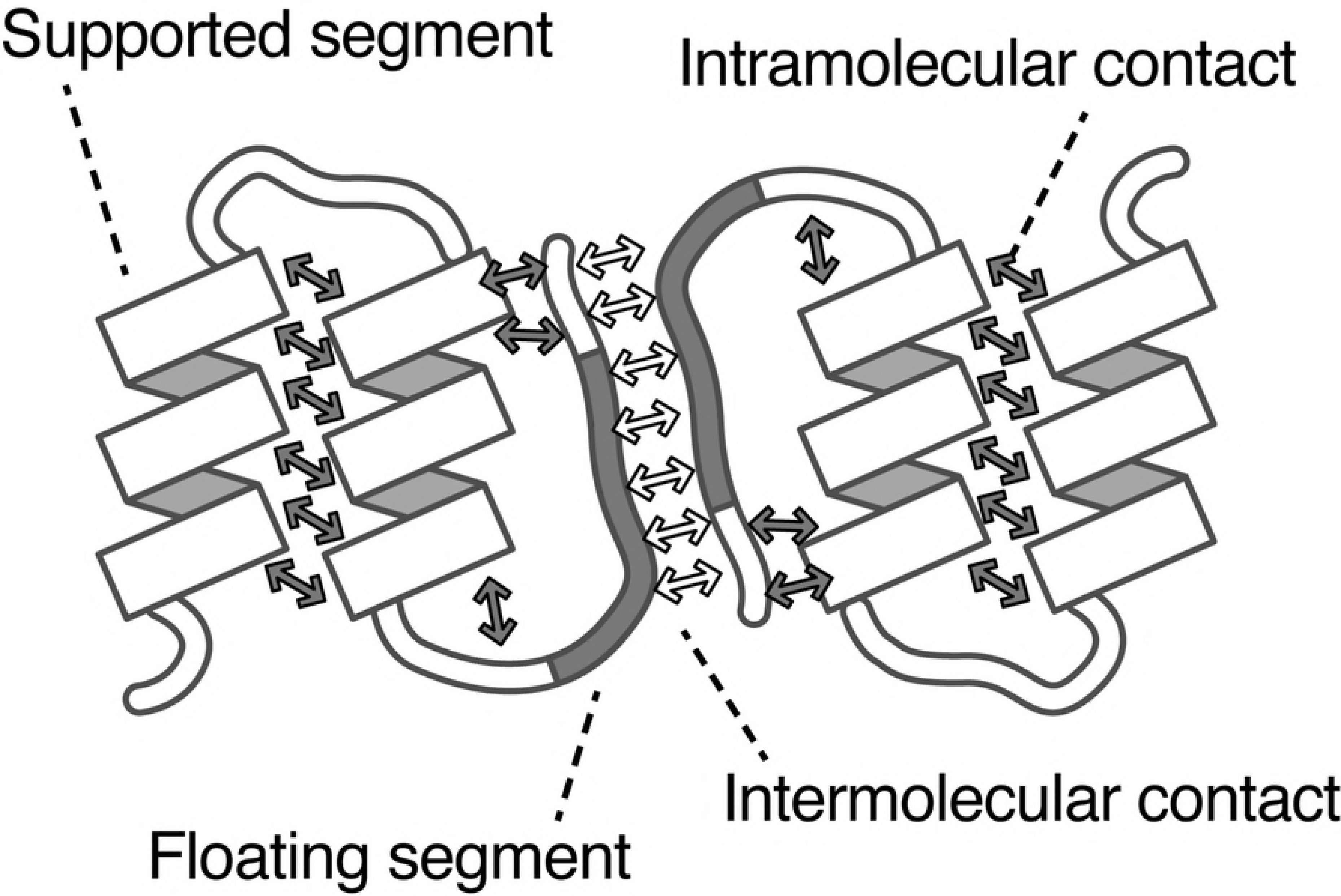
Schematic illustration of *floating* and *supported* segments. Open and shaded arrows indicate intermolecular and intramolecular contacts, respectively. The shaded regions in polypeptides are floating segments.

## MATERIALS AND METHODS

### Dataset Construction

In this study, we constructed two kinds of datasets, named *primary dataset* and *non-redundant datasets*. The latter are subsets of the first. The primary dataset was constructed by extracting entries from a snapshot of the PDB on June 14, 2017, with the following criteria: (i) the entry contains at least one polypeptide, (ii) the number of atoms is less than one million, and (iii) the structure was solved with X-ray crystallography with resolution better than, or equal to, 3.0 Å. Extracting information from the PDB was performed by parsing PDBML [14] with in-house scripts (S1 Data).

Non-redundant datasets were constructed by picking non-redundant entries from the primary dataset. Single-linkage clustering with 40 % sequence identity was performed using the CD-HIT program [15]. In cases where an entry had more than one chain, the sequence identity of the most similar pair of chains was considered. A non-redundant dataset was constructed by random picking of one entry from each cluster. We constructed 100 non-redundant datasets with different random seeds, and statistics involving them were analyzed.

### Detecting Atomic Contacts and Definition of Structural Units

In this study, we assessed interatomic contacts between heavy atoms with the threshold that the interatomic distance is less than 5 Å. When a pair of amino acid residues has at least one interatomic contact, we considered that this pair of residues has an *inter-residue contact*, An inter-residue contact formed between residues of the same molecule is termed an *intramolecular contact*, which can be grouped into two classes: (i) non-local contact for the cases in which the two contacting residues are distant more than five residues in the sequence order, and (ii) local contact for other cases. On the other hand, an *intermolecular contact* is defined as a contact between two different molecules. On the basis of inter-residue contacts, we defined a structural unit named *segment*. A unit consisting of more than three successive amino acid residues with non-local contacts is defined as a *supported* segment, and that without non-local contacts is defined as a *floating* segment (Fig 1).

### Analyses

We characterized segments in polypeptide chains by focusing on the following points: (i) segment type, defined as floating or supported, (ii) segment length, (iii) secondary structural elements (SSEs), (iv) accessible surface area (ASA), and (v) types of inter-molecular contact partners. (i) The segment type is signified as *T*_*seg*_ ⊂ {*flo,sup*}; *flo* and *sup* mean floating and supported segments, respectively. (ii) Segment length *L*_*seg*_ is the number of consecutive amino acid residues composing the segment. For simplicity, segment lengths fell into three classes: *T*_*len*_ 2282 {*short,middle,long*}, that were defined as 2 < *L*_*seg*_ ≤ 4, 4 < *L*_*seg*_ ≤ 9, and 9 < *L*_*seg*_, respectively. (iii) The type of SSE (*T*_*SSE*_) was assessed by using the DSSP program [16]. In this manuscript, we applied three categories of SSE; *T*_*SSE*_ ⊂ {*helix,beta, coil*}; *helix* was *G, H* or *I* in the DSSP classification, *beta* was *B, E, T*, or *S*, and *coil* was the others. The representative *T*_*SSE*_ of each segment was the most frequent SSE in the residues composing the segment. (iv) The solvent accessibility of each segment was defined as the two levels: surface and buried, *T*_*ASA*_ ⊂ {*sur, bur*}. For each amino acid residue, when the ratio of ASA of the residue to that of the same amino acid in Gly-X-Gly motifs is greater than 0.2, the residue is assumed to be surface exposed. Otherwise, it is assumed to be buried. ASA was calculated with the DSSP program [16]. (v) Partners of intermolecular contacts fell into three classes: polypeptides, polynucleotides, and small compounds. These were defined by the entity type as described in the PDB annotation. We considered non-polymer entities ≥ 300 Da as small compounds. The type of interaction partner of a segment is denoted as *T*_*int*_ ⊂ {*pep,nuc,sc*} for polypeptide, polynucleotide, and small compound, respectively.

Relative frequencies of various types of segments were analyzed. *F(x)* indicates the relative frequency of segments with the condition *x*,

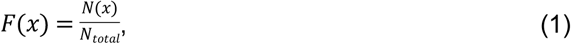

where *N*_*total*_ denotes the total number of segments in a dataset, and *N(x)* denotes the number of segments with the condition *x*. For example, the relative frequency of short segments and that of helix segments are represented as *F(T*_*len*_*=short)* and *F(T*_*SSE*_*=helix)*, respectively. For simplicity, they can also be denoted as *F(short)* and *F(helix)*. The conditional relative frequency, the ratio of the number of segments with types *x* and *y* to that with type *y*, is denoted as *F(x*/*y)*.

In addition, we also analyzed characteristics of residues. The relative frequency of residues with the condition *x* is presented as *F*^*res*^*(x)*. The amino acid type of each residue is denoted as 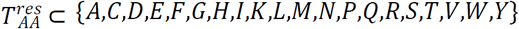. The propensity score of each amino acid was assessed with the log-odds score.*y*

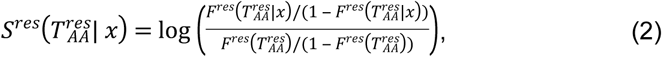

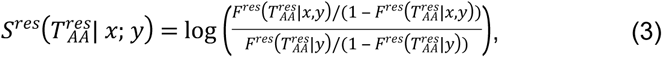

where *x* and *y* indicate conditions. We evaluated the amino acid propensities for segment types 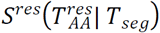, those for interaction partners in each segment type 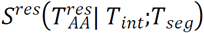, and those for surface or buried residues 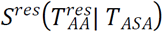.

## RESULTS AND DISCUSSION

### Database Statistics

The primary dataset consisted of 89,038 PDB entries, 115,287 entities, 225,366 chains, and 53,497,598 residues. Single-linkage clustering with 40 % sequence identity resulted in 15,351 clusters. Among these, 12,513 clusters (81.51 %) had less than, or equal to, five members, and 6,846 clusters (44.60 %) of these were singleton clusters (S1 Figure). A non-redundant dataset was constructed by randomly picking one entry from each cluster, and statistical analyses were performed on 100 non-redundant datasets. On average [with standard deviation (SD)], a non-redundant dataset consisted of 17,259.01 [37.44] entities, 38,601.25 [140.30] chains, and 8,967,882 [34,917.46] residues.

For the PDB entries, we defined floating segments as having at least three consecutive residues without non-local contacts, and defined supported segments as those with non-local contacts, respectively. The primary dataset included 359,501 floating segments and 4,419,603 supported ones, with approximately half of residues belonging to a segment. The average numbers [and SD] of floating and supported segments in the non-redundant datasets were 64,285.16 [387.25] and 725,750.9 [2,960.58], respectively. 0.72 % of residues were in floating segments.

### Segment Length and Secondary Structure Elements

Distributions of the segment length *L*_*seg*_ for each segment type are shown in Fig 2. The number of segments decreases exponentially, along with an increase in the segment length. In particular, there is a steep decrease in the frequency for floating segments in *L*_*seg*_ < 9. A majority of floating segments consist of only three or four residues (*F(short*/*flo)* = 0.850), and the ratio of floating segments that are longer than nine residues is only *F(long*/*flo)* = 0.014. On the other hand, many supported segments have more than nine segments (*F(long*/*sup)* = 0.103; *F(short*/*sup)* = 0.568). Floating segments tend to be shorter than supported ones. This may reflect the fact that longer protein regions without molecular contacts should have higher flexibility, and thus it is difficult to identify atomic coordinates of such regions by crystallography. The long flexible regions, such as intrinsically disordered regions, are abundant in nature although their atomic coordinates are not recorded in the PDB. Note that in this study we did not consider missing regions in the PDB entries, which means regions without atomic coordinates.

**Figure 2.**
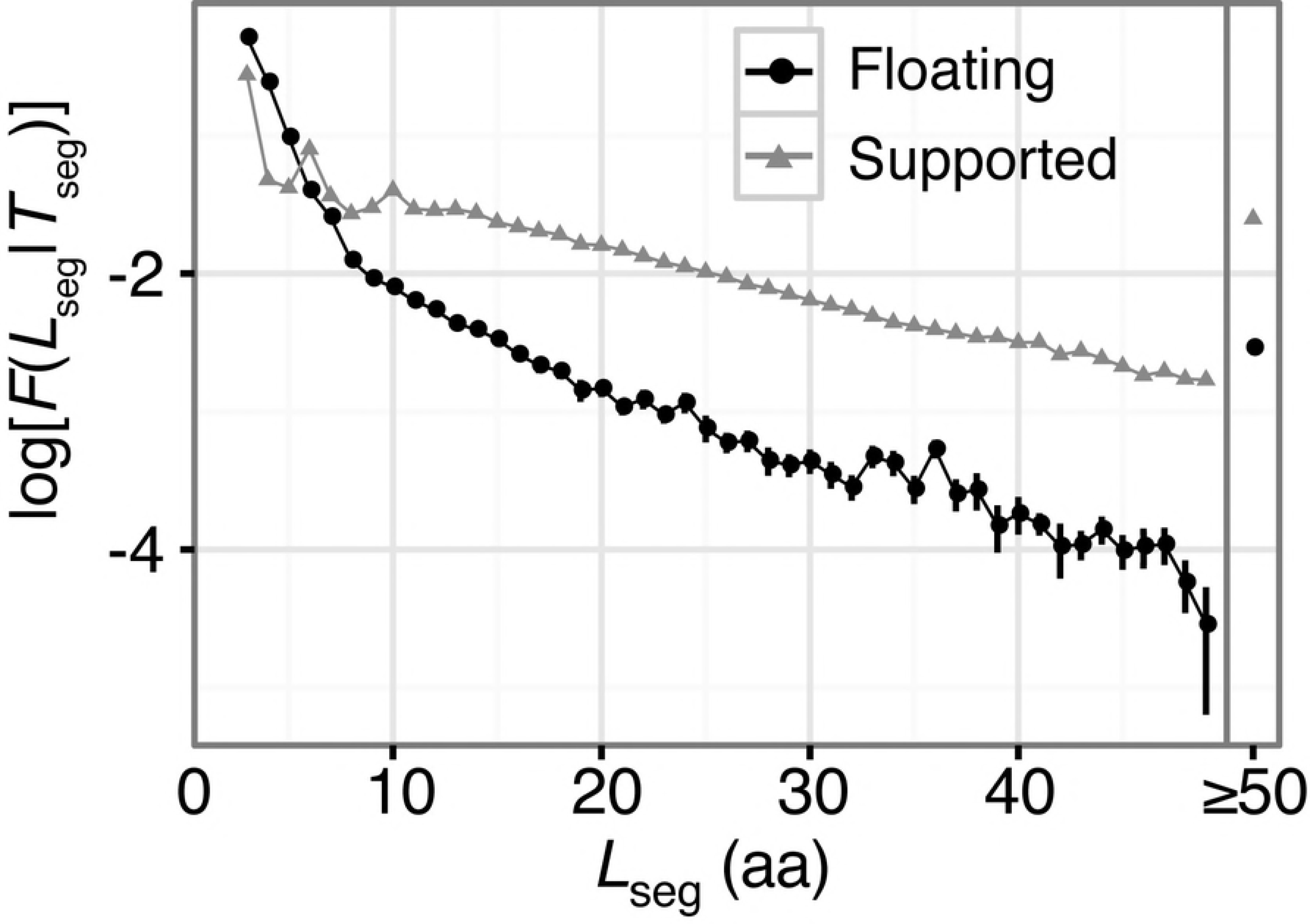
Histogram of floating (black) and supported (grey) segments regarding segment lengths. Error bars are the standard deviations in 100 non-redundant datasets.

Segment length relates to the secondary structure formation of segments. Longer segments tend to favor formation of helical structures, and only a small fraction of β-structures is observed in long floating segments (Fig 3). Namely, *F(helix*/*long, flo)* = 0.664, and *F(beta*/*long, flo)* = 0.0410. This result reflects the fact that a floating, single α-helix is more stable than a floating single β-strand. A majority of long floating β-strands in the dataset have intermolecular contacts. A typical case is formation of an intermolecular β-sheet (S2A Figure; PDB ID: 2O8M). On the other hand, floating helical segments are mainly observed at a solvated terminus of the chain (S2B Figure; PDB ID: 2WN9), in a coiled-coil (S2C Figure; PDB ID: 2GL2), and other types of interfaces (S2D Figure; PDB ID: 3RK0). For cases of unstructured (coil), long floating segments, typical cases are at the termini of proteins (S2E Figure; PDB ID: 3TER). The segments in the linker region are also observed (S2F Figure; PDB ID: 2B58). In contrast to the floating segments, the β-structures are not disfavored in long-supported segments (Fig 3). Since β-stands are usually formed as a part of a β-sheet, broad lengths of supported β-stands can be found in protein structures. Since the usual size of β-sheets accords to the category of *T*_*len*_ = *middle*, a higher ratio of beta structures in the middle-supported segments are observed, compared with other lengths. Some examples of supported segments in each length are shown in S3A, B, and C Figures. Regarding coil structures, in contrast to the case of floating segments, only a small ratio of longer-supported segments adopt coiled structures. Typical long-supported segments with coil structures penetrate inside the domain (S3D Figure; PDB ID: 2BSL), or locate around the surface of the domain (S3E Figure; PDB ID: 3TEH).

**Figure 3.**
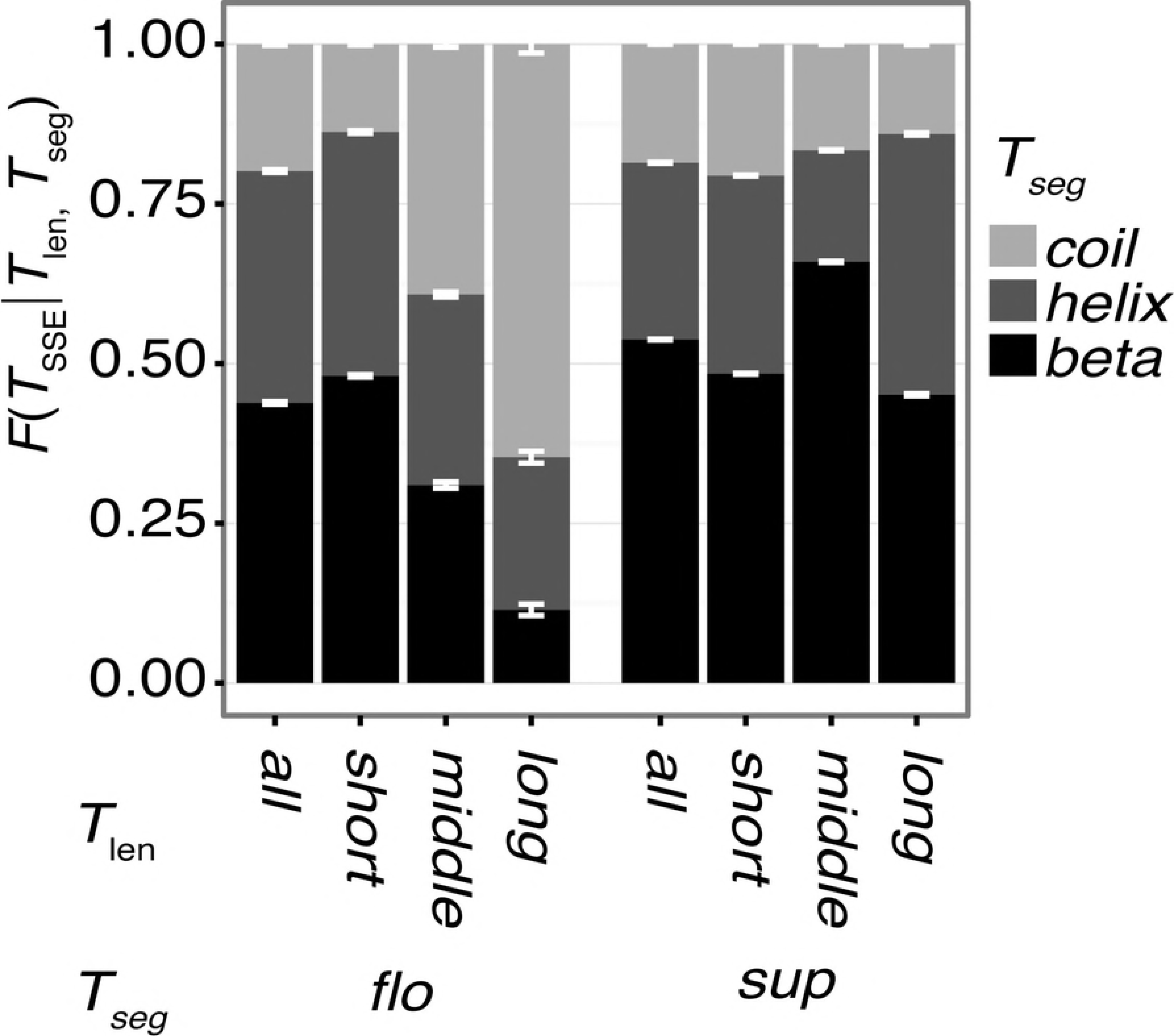
Ratio of the SSE for each combination of classes: *T*_len_ and *T*_seg_. Black, dark grey, and light grey denote the ratios of *beta, helix*, and *coil* structures, respectively. Error bars are the standard deviations in 100 non-redundant datasets.

### Segments as interfaces of intermolecular interactions

The relative frequencies of segments for intermolecular interactions are summarized in Fig 4. The interaction partners were categorized into one of the three types; that is, polypeptides (*T*_*int*_ = *pep*), polynucleotides (*T*_*int*_ *= nuc)*, and small compounds (*T*_*int*_ *= sc)*. For interactions with polypeptides and polynucleotides, floating segments exhibit a higher frequency of appearance in intermolecular interfaces compared with supported segments; the ratio of interacting floating segments are *F(pep*/*flo) =* 0.424, *F(nuc*/*flo)* = 0.0121, and those values for supported segments are *F(pep*/*sup)* = 0.282 and *F(nuc*/*sup)* = 0.00659, respectively. This is because intermolecular interactions require high solvent accessibility, and almost all floating segments are on the surface (*F(sur*/*flo)* = 0.953) and this ratio in supported segments is *F(sur*/*sup)* = 0.397. However, although floating segments are enriched at the surface, binding sites for small compounds prefer supported segments rather than floating ones; the ratios of interacting-floating and interacting-supported segments are *F(sc*/*flo)* = 0.0190 and *F(sc*/*sup) =* 0.0710, respectively. Since the binding sites are usually formed as a concave surface (called a cavity or pocket) with a certain size and depth [17], they should be formed by supported segments rather than by floating ones. As an example of binding sites with floating segments, 3-chlorocatechol 1,2-dioxygenase binds its ligand with a floating helix-turn-helix conformation (S4A Figure; PDB ID: 2BOY). In addition, many entries in this category do not have biologically relevant ligand-binding sites but have contacts with other non-specific small molecules such as lipids. For instance, a light harvesting complex is surrounded by chlorophyll molecules (S4B Figure; PDB ID: 3PL9).

**Figure 4.**
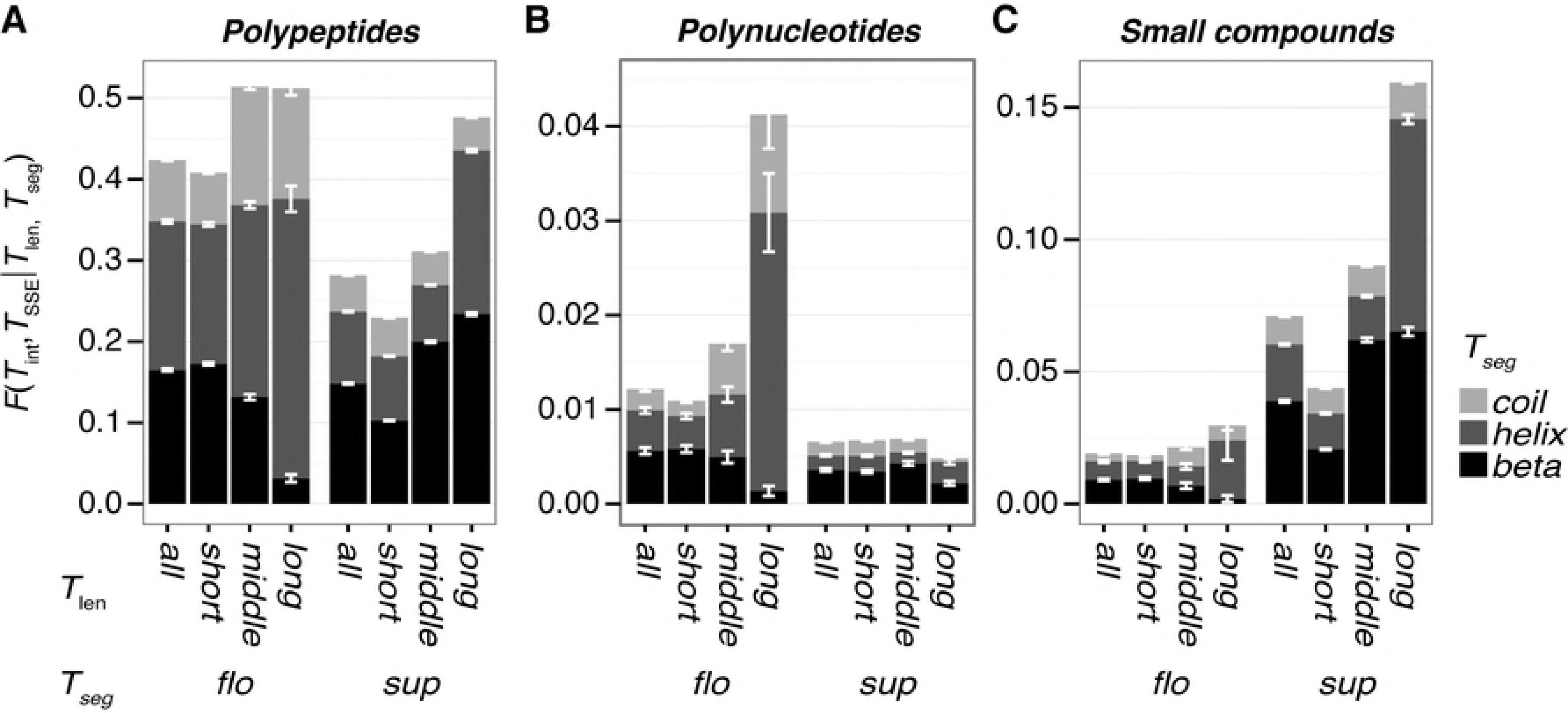
The ratios of interacting segments for each combination of classes: *T*_SSE_, *T*_len_ and *T*_seg_. (A) Ratios for interactions with polypeptides. (B) Ratios for interactions with polynucleotides. (C) Ratios for interactions with small compounds. Black, dark grey, and light grey denote the ratios of *beta, helix*, and *coil* structures, respectively. Error bars are the standard deviations in 100 non-redundant datasets.

Regarding the segment length, longer segments show higher frequencies for interactions since longer segments have larger surface areas, which simply elevates the probability of interactions. For floating segments, typical interactions of long helix and long beta segments are formation of coiled-coil structures and inter-molecular β-sheets, respectively, as shown above (S2C Figure and S2A Figure, respectively). As an exception to the tendency of the segment length, longer supported segments are not enriched in polynucleotide binding sites. Many typical double-stranded DNA binding sites include floating segments with positively charged amino acid residues. For example, bZIP heterodimeric complexes recognize DNA with two floating helices (S5A Figure; PDB ID: 2WT7). A floating-helix segment consisting of 18 residues in the NC2–TBP–DNA ternary complex structure recognizes the DNA with their six acidic residues (S5B Figure; PDB ID: 1JFI). A linker loop with two acidic residues in a replication terminator protein is buried into the major groove of DNA (S5C Figure; PDB ID: 1ECR). In contrast, it is sterically difficult to attach grooves of a double-stranded DNA to supported segments. A majority of supported segments at the polynucleotide binding sites touch the DNA backbone rather than burying into the grooves (for example PDB ID: 3E3Y; S5D Figure). In addition, many single-stranded DNA binding sites are composed of supported segments (for example S5E and F Figures; PDB ID: 2KFN and 3CMW, respectively).

### Propensities of Amino Acids

Propensity of each amino acid for floating and supported segments was assessed based on the log-odds score (Fig 5B; Eq. 2). In general, bulky non-polar amino acids, e.g., Cys, Phe, Ile, Leu, Met, Val, Trp, and Tyr, are disfavored for floating segments. This tendency is similar for surface residues (Fig 6; the Pearson correlation coefficient (PCC) of propensity scores between floating segments and surface residues is 0.929). This is due to the fact that a majority of floating segments are at the surface; *F(sur*/*flo)* = 0.953. However, there are some unique features in the amino acid propensity for the floating segments compared to surface residues. (i) Arg and His are disfavored in floating segments, although they are highly enriched as surface residues due to their high polarity. In many cases, they are involved in interfaces of intermolecular contacts. Arg is enriched for contacts with polynucleotides. For example, a 20-residue segment in the Fos-Jun complex has seven Arg residues (S6A Figure; PDB ID: 1A02). At an interface with polypeptides, Arg can stabilize the interactions through formation of salt bridges (S6B Figure; PDB ID: 2E7S). Many His residues in floating segments are observed in His-tag sequences.(ii) Gly is preferred for floating segments, although it is not so favored in surface residues. This is due to its high flexibility, which makes it possible to form unstructured regions, including loops and linkers. For example, a 12-residue segment in a loop region of MHC molecules has five Gly residues (S6C Figure; PDB ID: 1LNU). (iii) Ala is not so disfavored for the floating segments, in spite of its negative score in surface residues due to its hydrophobic side chain. One possible explanation is that α-helices favor Ala residues [7,18]. Floating segments show higher ratios of helix conformation than supported segments (Fig 3). An example of a floating helix with many Ala residues is shown in S6D Figure (PDB ID: 4KE2).

We also assessed amino acid propensities for intermolecular interactions with the three categories of molecules; polypeptides, polynucleotides, and small molecules (Figs 5C, D, and E, respectively). The propensity score of floating segments for polypeptide interactions (black in Fig 5C) shows the opposite trend from that of propensity to form floating segments (Fig 6B; their PCC is - 0.872). Although hydrophobic amino acids are disfavored for floating segments, they are favored for intermolecular interaction interfaces with other polypeptides. This implies that when disfavored amino acids exist in a floating segment, it is expected that they conduct some functions to recognize another proteins. The exception is Gln, which has a positive propensity score for both conditions; *S*^*res*^*(Q;flo)* and *S*^*res*^*(Q*; *pep*/*flo)* shown in Figs 5B and C, respectively. Gln is often observed at terminal or kinking regions of a helix (S6E and F Figures for examples). The propensity for floating segments in small-molecule binding sites showed a weak correlation to that for polypeptides (Fig 6C; the PCC is 0.614). The major differences are as follows: Gly and Pro are favored, and Gln is disfavored for floating small-molecule binding sites. Since Gly and Pro are enriched in flexible regions, they are observed in loop regions composing a binding site (examples are shown in S6G Figure and H). For the propensity for floating segments in interfaces to polynucleotide, there is no clear correlation with other propensities. In a comparison with the trend in supported segments, floating segments disfavor some hydrophobic amino acids, e.g., Cys, Leu, Pro, and Trp. In addition, while Asp and Glu are disfavored, Asn and Gln are not. They sometimes have direct contacts with a base of polynucleotides (S6I Figure).

**Figure 5.**
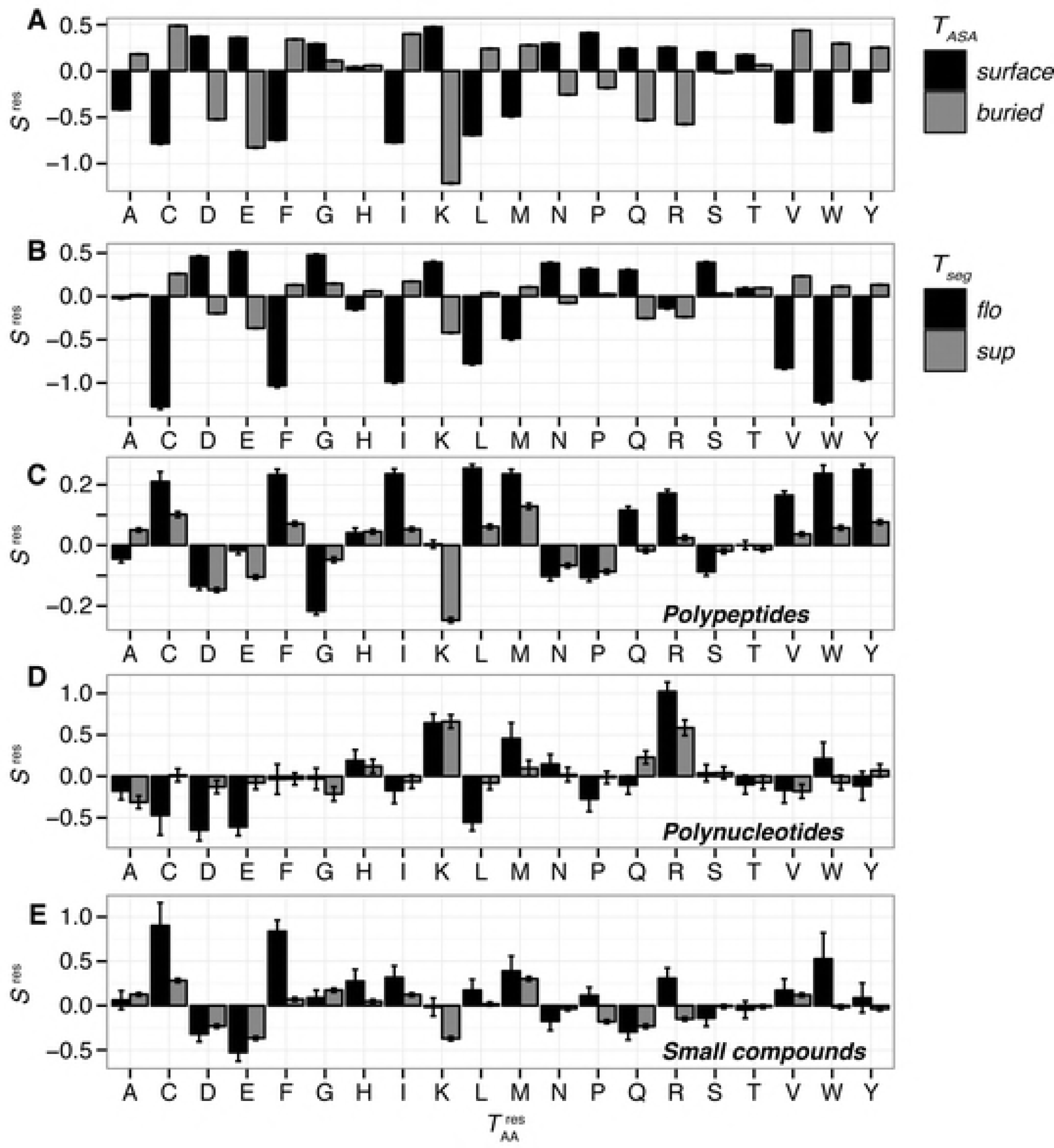
Amino acid propensity scores. (A) The propensities for surface or buried residues; 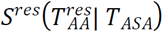. Black and grey bars denote surface and buried residues for the panel, respectively. (B) The propensities to form floating or supported segments; 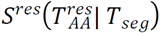. (C) The propensities to interact with polypeptides; 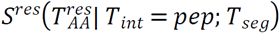. (D) The propensities to interact with polynucleotides; 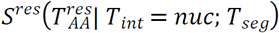. (E) The propensities to interact with small compounds; 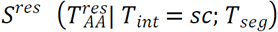. Black and grey bars indicate floating and supported segments for the panels (B), (C), (D), and (E). Error bars are the standard deviations in 100 non-redundant datasets.

**Figure 6.**
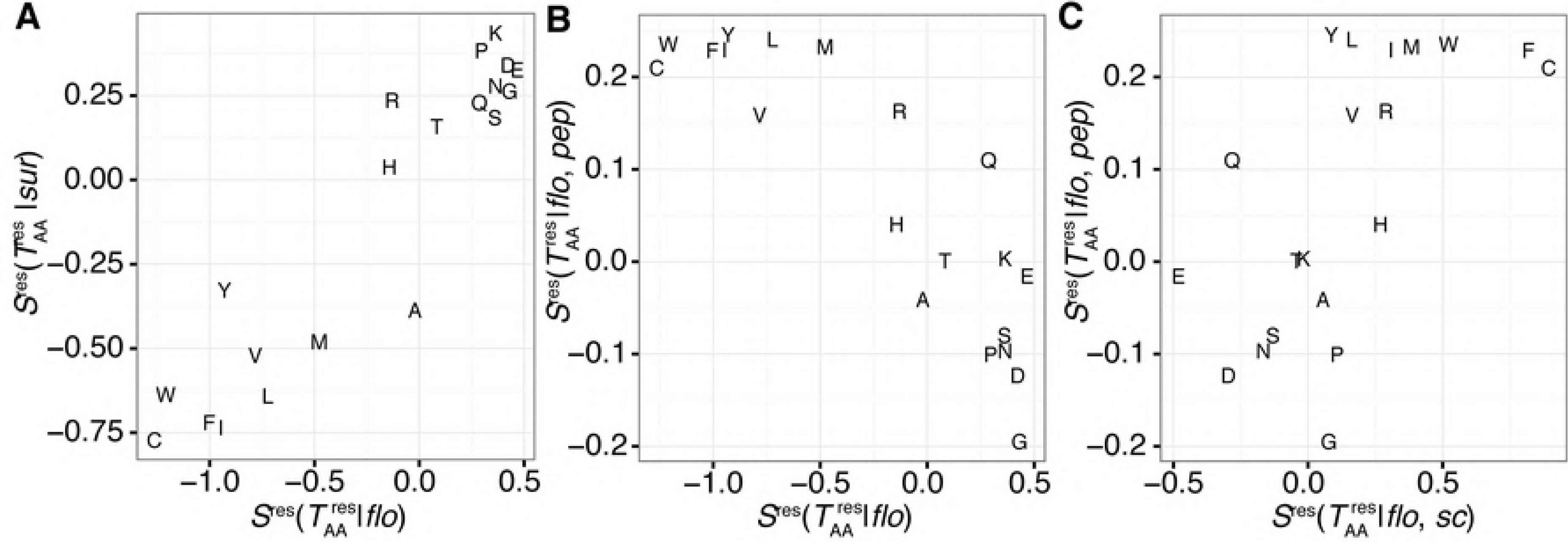
Comparisons of the amino acid propensity scores shown in Fig 5. (A) Comparison between floating and surface segments. (B) Comparison between floating segments and those with peptide interactions. (C) Comparison between floating segments with small compound interactions and those with peptide interactions.

## CONCLUSIONS

In this study, we defined floating and supported segments involved in the 3D structure of proteins (Fig 1), and characterized them on the basis of statistical analyses of the PDB. We found considerable numbers of floating segments in known protein structures (0.72 % of residues are in floating segments). The frequency distribution of segment length shows exponential decay along with an increase in the segment length, in both floating and supported segments. The length distribution of floating segments is more biased toward shorter regions than that of supported segments, and most of the floating segments are composed of three or four residues (Fig 2). Three is the minimum length of a segment in the definition; the segment length largely impacts its characteristics. Shorter floating segments tend to form secondary structures (Fig 3). Longer floating segments are enriched in intermolecular interaction interfaces. In particular, beta structures are favored for long-floating segments at the interfaces (Fig 4). Although floating segments are enriched at interfaces for polypeptides and polynucleotides, they are disfavored at interfaces for small compounds (Fig 4). Regarding the amino acid composition, while floating segments are basically similar to surface exposed residues, they have some unique features; higher preferences for small side chains (Gly and Ala) and disfavoring some charged side chains (Arg and His) compared to surface residues (Fig 5A). Interestingly, the propensity scores for polypeptide interactions of floating segments are in an opposite trend from that for all floating segments (Figs 5A and B). Residues disfavored for floating residues tend to be interfaces for protein–protein interactions at floating segments, except for Gln residues.

## ACKNOWLEDGEMENT

Supercomputer resources were provided by the National Institute of Genetics, Research Organization of Information and Systems, Japan.

---

**S1 Figure.** The number of PDB entries in each cluster.

**S2 Figure.** Examples of long floating segments. The target segment, the chain including the segment, and other chains are shown in red, green, and gray. (A) A floating-beta segment composing an intermolecular β-sheet (a virus serine protease; PDB ID: 2O8M). (B) A floating-helix segment at a solvated terminus (an acetylcholine receptor; PDB ID: 2WN9). (C) A floating-helix segment in a coiled-coil (adhesin FadA; PDB ID: 2GL2). (D) A floating-helix segment at an intermolecular interface (PDB ID: 3RKO). (E) A floating-coil segment at a terminus (the membrane domain of respiratory complex I; PDB ID: 3TER). (F) A floating-coil segment at a linker region (diamine acetyltransferase 1; PDB ID: 2B58).

**S3 Figure.** Examples of supported segments. The target segment, the chain including the segment, and other chains are shown in red, green, and gray. (A) A supported-short segment in a β-sheet (a putative nucleotide-diphospho-sugar transferase; PDB ID: 3CGX). (B) A supported-short segment in a β-sheet (kinase PhoQ catalytic domain; PDB ID: 3CGZ). (C) A supported-long segment in a β-sheet (a tRNA synthetase; PDB ID: 3TEG). (D) A supported-coil segment penetrating a globular domain (a dihydroorotate dehydrogenase A; PDB ID: 2BSL). (E) A supported-coil segment surrounding a globular domain (a tRNA synthetase; PDB ID: 3TEH).

**S4 Figure.** Examples of small-compound binding sites with floating segments. The target segment, the chain including the segment, and other chains are shown in red, green, and gray. (A) A floating segment at a small-compound binding site. (3-Chlorocatechol 1,2-Dioxygenase; PDB ID: 2BOY, (B) A floating segment interacting lipid molecules (a chlorophyll binding protein; PDB ID: 3PL9).

**S5 Figure.** Examples of nucleic acid binding sites with floating segments. The target segment, the chain including the segment, and other chains are shown in red, green, and gray. (A) A floating-helix segment in a bZIP heterodimeric complex (PDB ID: 2WT7). (B) A floating-helix segment in the NC2– TBP–DNA ternary complex (PDB ID: 1JFI). (C) A coil-floating segment in a replication terminator protein (PDB ID: 1ECR). (D) A supported segment in the restriction enzyme HindII (PDB ID: 3E3Y). (E) A supported segment in the Klenow fragment of a DNA polymerase (PDB ID: 2KFN). (F) A supported segment in RecA (PDB ID: 3CMW).

**S6 Figure.** Examples of floating segments with specific amino acids. The target amino acid residues, the segment including the residues, the chain including the segment are shown in cyan, red, and green, respectively. The binding partners are shown in yellow. (A) A floating segment with many Arg residues in Jun-Fos heterodimer (PDB ID: 1A02). This segment recognizes the double-stranded DNA. (B) A floating segment including Arg in the yeast Sec2p GEF domain (PDB ID: 2E7S). Arg forms the salt-bridge with Asp in the other chain. (C) A floating segment with Gly residues in a MHC molecule (PDB ID: 1LNU). (D) A floating segment with Ala residues in Type I hyperactive antifreeze protein (PDB ID: 4KE2). (E, F) Floating fragments with Gln residues interacting with the other polypeptide: (E) Huntingtin (PDB ID: 4FE8) and (F) enoyl reductase InhA (PDB ID: 4R9R). (G, H) Floating segments including Gly residues at the binding interface to the small molecule: (G) HIV-1 protease (PDB ID: 1SH9), and (H) ecdysone receptor (PDB ID: 2R40). (I) A floating segment interacting with the RNA by Pro residues in virus capsid (PDB ID: 1DDL). (J) A floating segment interacting with the siRNA duplex by Asn residues in Piwi protein (PDB ID: 2GBB).

**S1 Data.** Segments are recorded as a .csv file. Each column indicate PDB ID, the type of segment (supported or floating), the length of segment, the type of length (short, medium, or long), the secondary structural elements (helix, beta, or coil), the number of interacting molecules for polypeptide, polynucleotide, and small compound, and the sequence of the segment.

